# Revisiting australian *ectocarpus subulatus* (phaeophyceae) from the hopkins river: distribution, abiotic environment, and associated microbiota

**DOI:** 10.1101/821579

**Authors:** Simon M. Dittami, Akira F. Peters, John West, Thierry Cariou, Hetty KleinJan, Bertille Burgunter-Delamare, Aurelie Prechoux, Suhelen Egan, Catherine Boyen

## Abstract

*Ectocarpus* is a genus of common marine brown algae. In 1995 a strain of *Ectocarpus* was isolated from Hopkins River Falls, Victoria, Australia, constituting one of few available freshwater or nearly freshwater brown algae, and the only one belonging to *Ectocarpus*. It has since been used as a model to study acclimation and adaptation to low salinities and the role of its microbiota in these processes. However, little is known about the distribution of this strain or whether it represents a stable population. Furthermore, its microbiota may have been impacted by the long period of cultivation.

Twenty-two years after the original finding we searched for *Ectocarpus* in the Hopkins River and surrounding areas. We found individuals with ITS and *cox*1 sequences identical to the original isolate at three sites upstream of Hopkins River Falls, but none at the original isolation site. The osmolarity of the water at these sites ranged from 74-170 mOsmol, and it was rich in sulfate. The diversity of bacteria associated with the algae *in situ* was approximately one order of magnitude higher than in previous studies of the original laboratory culture, and 95 alga-associated bacterial strains were isolated from *E. subulatus* filaments on site. In particular, *Planctomycetes* were abundant *in situ* but rare in the laboratory-cultured strain.

Our results confirm that *E. subulatus* has stably colonized the Hopkins River, and the newly isolated algal and bacterial strains offer new possibilities to study the adaptation of *Ectocarpus* to low salinity and its interactions with its microbiome.

## Introduction

Brown algae (Phaeophyceae) form the dominant vegetation in the tidal and sub-tidal zone of rocky shores in temperate marine environments, but they are rarely found in fresh water (Dittami *et al.* 2017): while there are ca. 2,000 known species of marine brown algae, covering a large range of morphologies from small filamentous algae to large and morphologically complex kelp species, there is only a handful of known freshwater brown algae, all of them small and with simple morphology (crust-forming or filamentous). Among these freshwater brown algae the genus *Ectocarpus* has a unique position, because it corresponds to a predominantly marine genus, which, on two occasions, has been recorded also in rivers: One occurrence of *Ectocarpus crouaniorum* in a highly salt-contaminated section of the Werra river in Germany (Geissler 1983), and one occurrence of *Ectocarpus subulatus* (Peters *et al.* 2015) in a nearly freshwater habitat (salinity 1ppt) in the Hopkins River, Victoria, Australia (West and Kraft 1996).

The isolate from the latter site (Culture Collection of Algae and Protozoa accession 1310/196), constitutes a potential model system to study marine-freshwater transitions in brown algae. The species *E. subulatus* (Peters *et al.* 2015) is related to the genomic model species *Ectocarpus siliculosus* (Cock *et al.* 2010) and has been reported in highly variable environments with high levels of abiotic stressors, such temperature at Port Aransas, Texas, USA (Bolton 1983). More recently, the genome of *E. subulatus* has been sequenced, revealing that *E. subulatu*s, in comparison to *Ectocarpus siliculosus*, has lost members of gene families down-regulated in low salinities, and conserved those that were up-regulated (Dittami *et al.* 2018, preprint). The *E. subulatus* strain from Hopkins River Falls has further been used for physiological experiments: it can grow in both seawater and fresh water and its transcriptomic and metabolic acclimation to these conditions has been examined (Dittami *et al.* 2012) along with the composition of its cell wall with regard to sulfated polysaccharides (Torode *et al.* 2015). Moreover, the capacity of the freshwater strain to grow in low salinities has been shown to depend on its associated microbial community (Dittami *et al.* 2016), and extensive efforts have been made to develop a collection of cultivable bacteria to study this phenomenon (KleinJan *et al.* 2017).

Despite this increasing quantity of data on the physiology of the Hopkins River Falls strain of *E. subulatus*, we currently know little about its ecology. The original paper describing its isolation (West and Kraft 1996) states that it was isolated on March 24^th^, 1995 from cracks between the basalt rock of the Hopkins River, just above the Hopkins River Falls. Water temperature was 16°C, salinity 1 ppt, and conductivity 3 mS·s^-1^. However, it is unknown if a stable population of *E. subulatus* is currently present at Hopkins River Falls, nor how far this population may extend. Furthermore, the culture has undergone > 20 years of cultivation in different laboratories, potentially having a strong impact on its associated microbiota.

In this study, we address both of these knowledge gaps by returning to the Hopkins River and searching for extant populations of this alga for the first time since its discovery 20 years ago. We found *E. subulatus* individuals at three locations along the Hopkins River, affirming the existence of a stable population at this site and isolating several novel alga-associated bacterial strains from these samples.

## Materials and methods

### Biological samples

The sampling campaign was carried out from March 21^st^ to March 27^th^, 2017 and covered selected locations along the Hopkins River between Warrnambool and Ararat (sites 1-15), as well as several sites along the Southern Australian Coastline between Port Fairy and Avalon (Figure 1, Table 1). If filamentous algae resembling *Ectocarpales* were present at a sampling site, small amounts of live samples were taken and rinsed three times in 0.2μm-filtered local water (3 replicates). A small piece of each sample was stored at max. 20°C in 2ml Eppendorf tubes filled with the surrounding water for live algal cultures. The second part of the samples was ground on-site according to Tapia *et al.* (2016), with 50 μL of 0.2μm-filtered local water in a sterile mortar and in the proximity of a Bunsen burner. One, seven, and 35 μL of the ground alga were diluted with 0.2μm-filtered local water to a final volume of 50μL and spread immediately onto pre-prepared R2A agar plates (Sigma-Aldrich, St. Louis, MO, USA) for isolation of culturable bacteria. These plates were kept at ambient temperature (max. 25°C) and were monitored for two weeks. Newly emerging colonies were purified once more on fresh R2A plates and then put into culture in liquid Zobell medium (Zobell 1941) with 8-fold reduced salt concentration, identified by 16S rRNA gene sequencing (see below), and put into stock at −80°C in 40% Glycerol. The remaining sample was dried using silica gel for downstream analysis of the microbial community composition, and frozen at −20°C after the sampling campaign.

**Table 1:** Overview of samples taken and species identified.

| Date | Location | Material sampled | *Ectocarpus* tissue found | GPS coordinates | Germling emergence | Sequence accession(s) |
| --- | --- | --- | --- | --- | --- | --- |
| 2017-03-22 | 1. Merri Island | pebbles/mussels in tide pool | no | -38.402116, 142.471548 | - | - |
| 2017-03-22 | 2. Point Ritchie | sand/ sandstone | no | -38.401275, 142.509318 | *Ectocarpus siliculosus* | LR735221 (*cox1*)<br>LR735414 (ITS) |
| 2017-03-22 | 3. Mahoney Road | sand, wood/plastic | no | -38.392103, 142.531644 | - | - |
| 2017-03-22 | 4. Smith Lane | mud, wood | no | -38.397984, 142.578286 | - | - |
| 2017-03-22 | 5. Allan's Ford | rock (granite) | no | -38.385325, 142.587398 | - | - |
| 2017-03-22 | 6. Donovans Lodge | granit, clay | no | -38.355110, 142.599250 | - | - |
| 2017-03-22 | 7. Hopkins River Falls | volcanic rock | no | -38.333509, 142.621352 | - | - |
| 2017-03-23 | 8. Framlingham Forest Reserve | rock, pebbles | *E. subulatus* Au8 (abundant) | -38.297064, 142.668291 | - | LR735222 (*cox1*)<br>LR735415 (ITS) |
| 2017-03-23 | 9. Kent's Ford | rock, pebbles | *E. subulatus* Au9 (abundant) | -38.191574, 142.698058 | - | LR735223 (*cox1*)<br>LR735416 (ITS) |
| 2017-03-23 | 10. Hexham | mud, wood | *E. subulatus* Au10 (rare) | -37.995732, 142.689141 | - | LR735224 (*cox1*)<br>LR735417 (ITS) |
| 2017-03-23 | 11. Chatsworth | mud, sand, detritus | no | -37.856357, 142.650644 | - | - |
| 2017-03-23 | 12. Wickliffe | sand, detritus | no | -37.694348, 142.726074 | - | - |
| 2017-03-24 | 13. Rossbridge | rock, pebbles | no | -37.480217, 142.849465 | - | - |
| 2017-03-23 | 14. Ararat | sand | no | -37.300211, 142.973979 | - | - |
| 2017-03-24 | 15. Green Hill Lake | pebbles | no | -37.295320, 142.979170 | - | - |
| 2017-03-24 | 16. Merri River, Warnambool | concrete | no | -38.362077, 142.484414 | - | - |
| 2017-03-24 | 17. Killarney beach | sand, lava pebbles, shells | no | -38.357966, 142.306880 | *Kuckuckia* sp. | LR735225 (*cox1*)<br>LR735418 (ITS) |
| 2017-03-25 | 18. Belfast Lough airport | sand, wood pollar | no | -38.361889, 142.262045 | - | - |
| 2017-03-25 | 19. Curdies River | rock | no | -38.519965, 142.833558 | - | - |
| 2017-03-25 | 20. Port Campbell | tide pool. sand | no | -38.620681, 142.992981 | *Feldmannia* sp. | LR735226 (*cox1*) |
| 2017-03-25 | 21. Gellibrand River | mud, detritus | no | -38.727482, 143.250932 | - | - |
| 2017-03-25 | 22. Aire River | mud, pebbles | no | -38.763797, 143.474727 | - | - |
| 2017-03-25 | 23. Wild Dog Creek | sand | no | -38.735911, 143.683545 | - | - |
| 2017-03-25 | 24. Smythe's Creek | pebbles, biofilm | no | -38.704648, 143.762856 | - | - |
| 2017-03-26 | 25. Lorne | tide pool, sand shells | no | -38.531281, 143.980994 | *Acinetospora* sp. | LR735227 (*cox1*) |
| 2017-03-26 | 26. Hovell's Creek | clay | no | -38.018825, 144.402156 | - | - |

**Figure 1:**
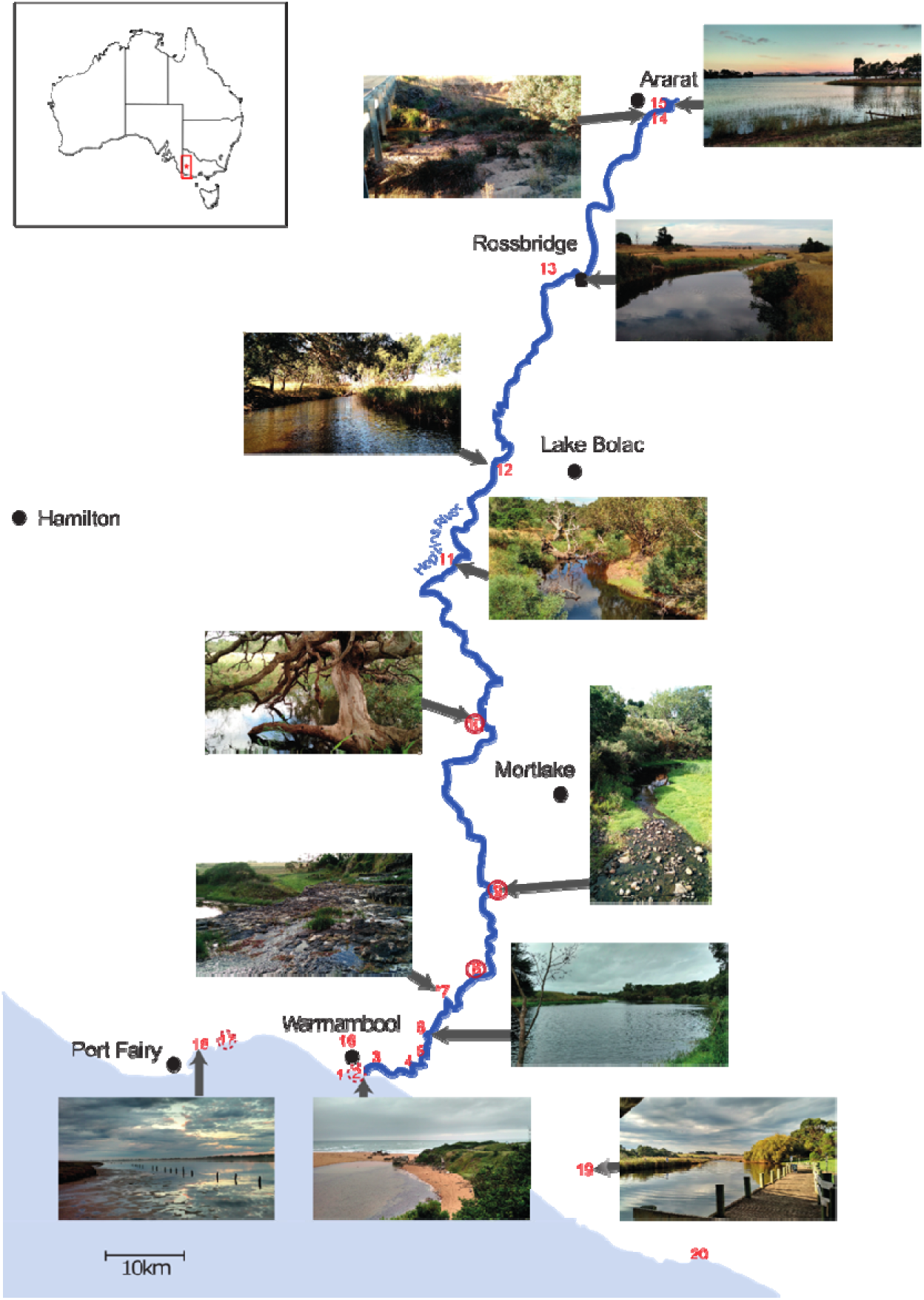
Sites sampled along the Hopkins River and nearby. Each sampling site is numbered (1-21). Solid circles indicate sites with *Ectocarpus* individuals (sites 8, 9, 10). Dotted circles (sites 2, 17) indicate site with Ectocarpales growing in germling emergence experiments. Hopkins River Falls, the original site of isolation, is located at site 7. The red box marked with * in the insert on the top left shows the position of the Hopkins River at the scale of the continent.

For all sites, we collected 3-7 sediment samples including small pieces of solid substrate (shells, pebbles, branches) if present. Approximately 0.1ml of sediment were kept as live samples in 2mL Eppendorf tubes. After two weeks these samples were transferred to fresh Provasoli-enriched (Starr and Zeikus 1993) medium based on 5%, 25%, or 100% seawater, depending on the osmolarity of the water at the sampling site. These sediment samples were then kept at 13°C in a 14/10 light-dark cycle at an irradiance level of 25 μmol PAR·m^-2^·s^-1^, and the emergence of *Ectocarpus*-like germlings was monitored over four months.

Both live algae collected *in situ* and those recovered from germling emergence experiments were cleaned by rigorous pipetting with a Pasteur pipette and several transfers to fresh, sterile, medium. Any diatoms that remained attached to the algal filaments were removed via treatment with 3mg·L^-1^ GeO_2_ for 3 weeks.

### Water samples

Approximately 100 ml of water were taken from each site, immediately filtered with 0.45 μM syringe filters to remove particulate matter, and then pasteurized for 1h at 95-100°C to remove any remaining bacterial activity. Filtered samples were then stored at ambient temperature until the end of the sampling campaign (max. 2 weeks) and then frozen at −20° C until analysis (ca. 3 months). The osmorality of water samples was determined using a Type 6 Osmomenter (Löser Messtechnik, Berlin, Germany). Phosphate, nitrite, and nitrate concentrations were determined using an AA3 auto-analyser (SEAL Analytical, Southampton, UK) following the method of Aminot and Kérouel (2007) with an accuracy of 0.02 μmol·L^-1^, 0.01 μmol·L^-1^, and 0.01 μmol L^-1^ for NO_3_^-^, NO_2_^-^, PO_4_^3-^, respectively. Sulfate concentrations were determined by high-performance anion-exchange chromatography (HPAEC), according to a protocol adapted from Préchoux *et al.* (2016). After suitable dilution, water samples were injected onto an IonPac™ AS11-HC column (4 × 250 mm) equipped with an AG11-HC guard column (4 × 50 mm), using an ICS-5000 Dionex system (SP-5 & Analytical CD Detector, Thermo Fisher Scientific, Waltham, MA, USA). Elution was performed with isocratic 12mM NaOH at a flow rate of 1 mL·min^-1^, and sulfate ions was detected in conductimetry mode (ASRS 500, 4 mm) and quantified using a standard calibration curve.

### Barcoding of algal and bacterial isolates

Algal isolates were identified using the mitochondrial *cox*1 and the nuclear ITS1+2 markers. Algal DNA was extracted from the cleaned cultures using the Macherey Nagel (Düren, Germany) NucleoSpin Plant II kit according to the manufacturer’s instructions (PL1 protocol with two 25 μL elutions), and 1μL of DNA (10-30 ng) was used in subsequent PCRs. For the ITS region, we used the AFP4LF (3’- CAATTATTGATCTTGAACGAGG-5’) and LSU38R (5’- CGCTTATTGATATGCTTA-3’) primers (Lundholm *et al.* 2003; Peters *et al.* 2004), and for the 5’*cox*1gene the GAZF2 (3’-CCAACCAYAAAGATATWGGTAC-5’) and GAZR2 (3’- GGATGACCAAARAACCAAAA-5’) primers (Lane *et al.* 2007), each at a final concentration of 0.5 μM. PCRs were carried out using a GoTaq polymerase and the following program: 2 min. 95 °C followed by 30 cycles [1 min 95 °C; 30 sec. 50°C for ITS or 55°C for *cox*1; 3 min 72 °C] and a final extension of 5 min 72 °C.

Bacterial cultures were identified by partial sequencing of their 16S rRNA gene. Fifty μL of dense bacterial culture were heated to 95°C for 15 min, spun down for 1 min, and 1 μL of supernatant was used as a template in a PCR reaction with the 8F (5’- AGAGTTTGATCCTGGCTCAG-3’) and 1492R 5’-GGTTACCTTGTTACGACTT-3’) (Weisburg *et al.* 1991) at a final concentration of 0.5 μM. Except for the annealing temperature (53°C here), the same PCR protocol as above was employed.

All PCR products were purified using ExoStar (Thermo Fisher Scientific) and the purified 16S rRNA gene amplicons were sequenced with Sanger technology at the GENOMER platform (FR2424, Roscoff Biological Station), using the BigDye Xterminator v3.1 cycle sequencing kit (Applied Biosystems, Waltham, MA, USA). For bacterial strains, sequencing was carried out only in one direction using the 8F primer, and for algal sequences both the forward and the reverse strand were sequenced and manually assembled. Sequence identification was carried out using RDP classifier (Wang *et al.* 2007) for bacterial 16S rRNA gene sequences, and BLAST searches against the NCBI nt database (July 2017) for algal sequences. They were further aligned together with reference sequences from the NCBI nt database using the MAFFT server (Katoh *et al.* 2002) and the G-INS-i algorithm. All positions with less than 95% site coverage were eliminated. Phylogenetic analyses were carried out with MEGA 7 (Kumar *et al.* 2016) using the Maximum Likelihood method based on the GTR+G+I model and 1,000 bootstrap replicates.

### Metabarcoding

Sufficient material for metabarcoding analyses was harvested at two of the three sites with *Ectocarpus* individuals: sites 8 and 9 (Figure 1). Approximately 20mg dry weight for each of the three replicate samples for each site were ground twice for 45 sec. at 30 Hz in a TissueLyser II (Qiagen, Hilden, Germany). DNA was then extracted using the Qiagen DNeasy Plant mini kit according to the manufacturer’s instructions. Approximately 50 ng of DNA (as estimated using a NanodropONE, Thermo Fisher Scientific), were then used to amplify the V3-V4 region of the 16S rRNA gene. Furthermore, a mock community comprising 26 bacterial genera (Thomas *et al.* in prep) as well as a negative control, were added alongside the samples. PCR amplification, indexing, and library construction were carried out following the standard “16S Metagenomic Sequencing Library Preparation” protocol (Part # 15044223 Rev. B). Final library concentrations were measured using a BioAnalyzer (Agilent, Santa Clara, CA, USA) before pooling. Libraries for each sample were then pooled in an equimolar way, diluted to 5nM final concentration and supplemented with 20%PhiX to add sufficient diversity for sequencing on an Illumina MiSeq using a 2 × 300bp cartridge. Raw data were deposited at the ENA under project accession number PRJEB34906.

Raw reads were first trimmed and filtered using the fastx_quality_trimmer script (http://hannonlab.cshl.edu/fastx_toolkit/), assembled using Pandaseq 2.11 (Masella *et al.* 2012) and further processed with mother according to the Miseq SOP (version April 4^th^, 2018). Sequences were aligned to the non-redundant SSU ref database version 132, chimeric sequences removed using Vsearch (Rognes *et al.* 2016), and operational taxonomic units (OTUs) defined based on a 97% identity threshold. Rare sequences (<5 reads across all samples) were removed from the final analyses. Taxonomic assignments were generated for both the raw reads and the final OTUs using the RDP classifier method (Wang *et al.* 2007). Non-metric multidimensional scaling (NMDS) of the OTU matrix was carried out in R 3.5.1 using the isoMDS function of the Vegan package and Bray-Curtis dissimilarity as a distance measure. An Analysis of Similarity (ANOSIM) was used to test for difference between the two sites (719 permutations). Alpha diversity was estimated using the Shannon index with e as a base and the diversity() function of the VEGAN package. Statistical differences between the two sites at the OTU level were assessed by t-tests on log-transformed abundance data with subsequent correction for multiple testing according to Benjamini and Hochberg (1995).

## Results

### Distribution of Ectocarpus subulatus along the Hopkins River

Despite extensive searches, no traces of *Ectocarpus* were found at the original isolation site of *E. subulatus* at Hopkins River Falls (site 7, Figure 1). *Ectocarpus* was, however, the dominant vegetation at two sites (Framlingham Forest reserve and Kent’s Ford, sites 8 and 9, Figure 1), which were approximately 12km (7mi) and 37 km (23 mi) upstream of Hopkins River Falls. The third finding of *Ectocarpus* was registered 83 km (51 mi) upstream (site 10), although only a few filaments were found at this site. The *cox*1 and ITS sequences obtained from *Ectocarpus* cultures from all three sites were identical to those of the strain isolated from Hopkins River Falls in 1995 (Figure 2). We found no *E. subulatus* individuals in neighboring rivers, along the coastline, or in germling emergence experiments, but we recorded a few emerging germlings of other Ectocarpales for marine samples (Table 1).

**Figure 2:**
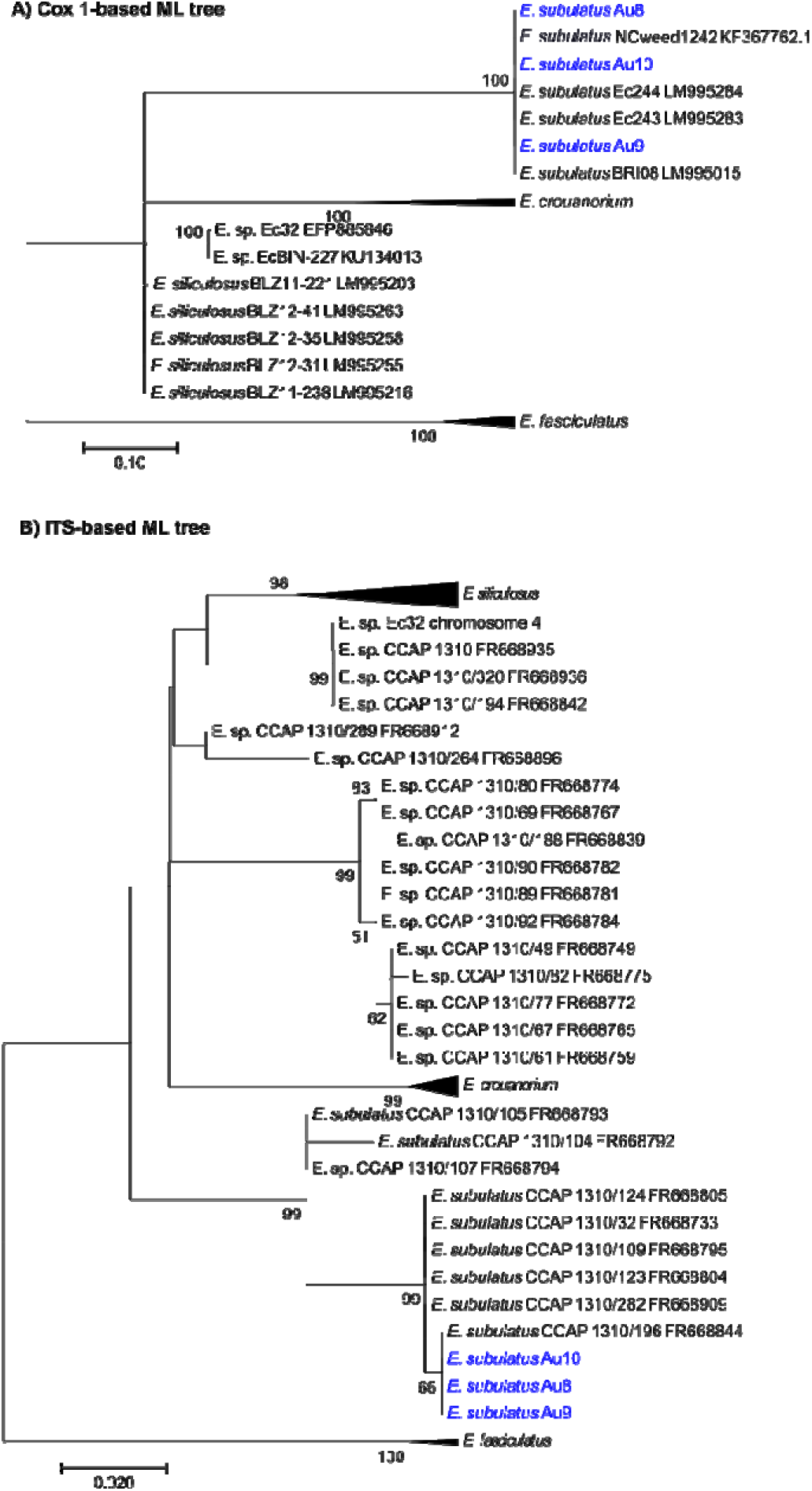
Maximum-likelihood tree of *Ectocarpus* isolates from Hopkins River and related strains. Panel A displays a tree based on the *COX1* gene (alignment of 677 bp after curation), and Panel B on the ITS region (860bp after curation). Blue color indicates isolates from this study. Support values correspond the percentage of support using 1000 bootstrap replicates.

### Water chemistry

The osmolarity of the Hopkins River, *i.e.* the total concentration of dissolved compounds, was highest close to the source (259 mOsmol; approximately ¼ that of seawater), and then gradually decreased towards the mouth of the river, where it dropped to ca. 40 mOsmol, before re-spiking due to the influence of seawater (Figure 3). This decrease corresponded to an increase in the flow of water masses, the Hopkins River constituting a nearly dried up creek at its source, which gradually grew into a river (see photos Figure 1). At the sites with *E. subulatus* individuals we estimated the speed of the current to range between 0.1 m·s^-1^ (10. Hexham) and 0.5 m·s^-1^ (8. Framlingham Forest). Sulfate concentrations followed the same pattern as osmolarity measurements and decreased from nearly 7mM to approximately 0.4mM close to the river mouth. Finally, phosphate and nitrite/nitrate concentrations were variable along the river, but especially low where *Ectocarpus* was found (Figure 3).

**Figure 3:**
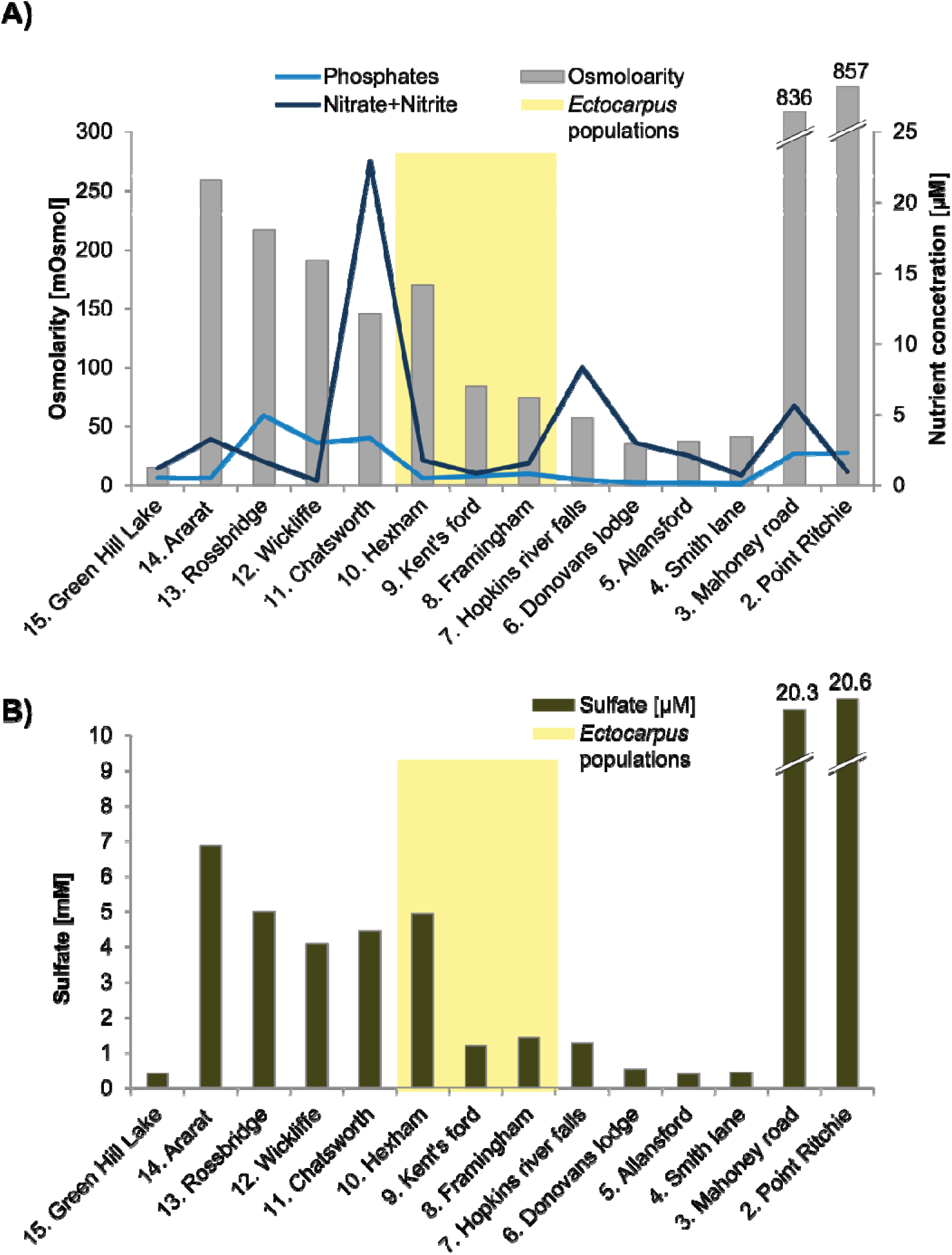
Water chemistry at the different sampling sites along the Hopkins River. (see Figure 1). Panel A displays osmolarity (gray bars) and nutrient concentrations (blue lines), and Panel B shows sulfate concentrations. Yellow background indicates sites with the occurrence of *Ectocarpus*.

### Bacterial communities associated with algae

*In situ* bacterial community composition was determined by 16S rRNA gene amplicon sequencing for the Framlingham Forest reserve (site 8) and Kent’s Ford (site 9) samples (Figure 4a, 4b). We detected 1312 OTUs across the three sampled individuals from both sites (Supporting Information File S1). Community composition between the two sites was overall similar (86% of OTUs, i.e. 1126, were without significant difference; Supporting Information File S1), and communities of both sites were dominated by *Alphaproteobacteria* (25% of reads), *Bacteriodetes* (20%), *Gammaproteobacteria* (8%), *Planctomycetes* (8%), and *Actinobacteria* (8%) (Figure 4a,b). Overall, the differences between sites were nevertheless statistically significant (ANOSIM p=0.001, Figure 4c). Notably, 86 OTUs were specific to Framlingham Forest reserve, and 60 more had a higher relative abundance there. At Kent’s Ford 27 OTUs were site-specific, and 13 more exhibited higher relative abundance (Supporting Information File S1). Alpha-diversity was also slightly higher at Framlingham Forest reserve (t-test p= 0.03; Figure 4d). Amplicon sequencing analyses were complemented by *in situ* isolation of bacterial strains from the algae after thorough rinsing with sterile river water (Figure 5). They comprise *Gammaproteobacteria* (48 isolates, including 28 *Pseudomonas*), *Firmicutes* (27 isolates), *Actinobacteria* (8 isolates), *Alphaproteobacteria* (7 isolates), and *Bacteriodetes* (5 isolates). No members of the *Planctomycetes* were isolated.

**Figure 4:**
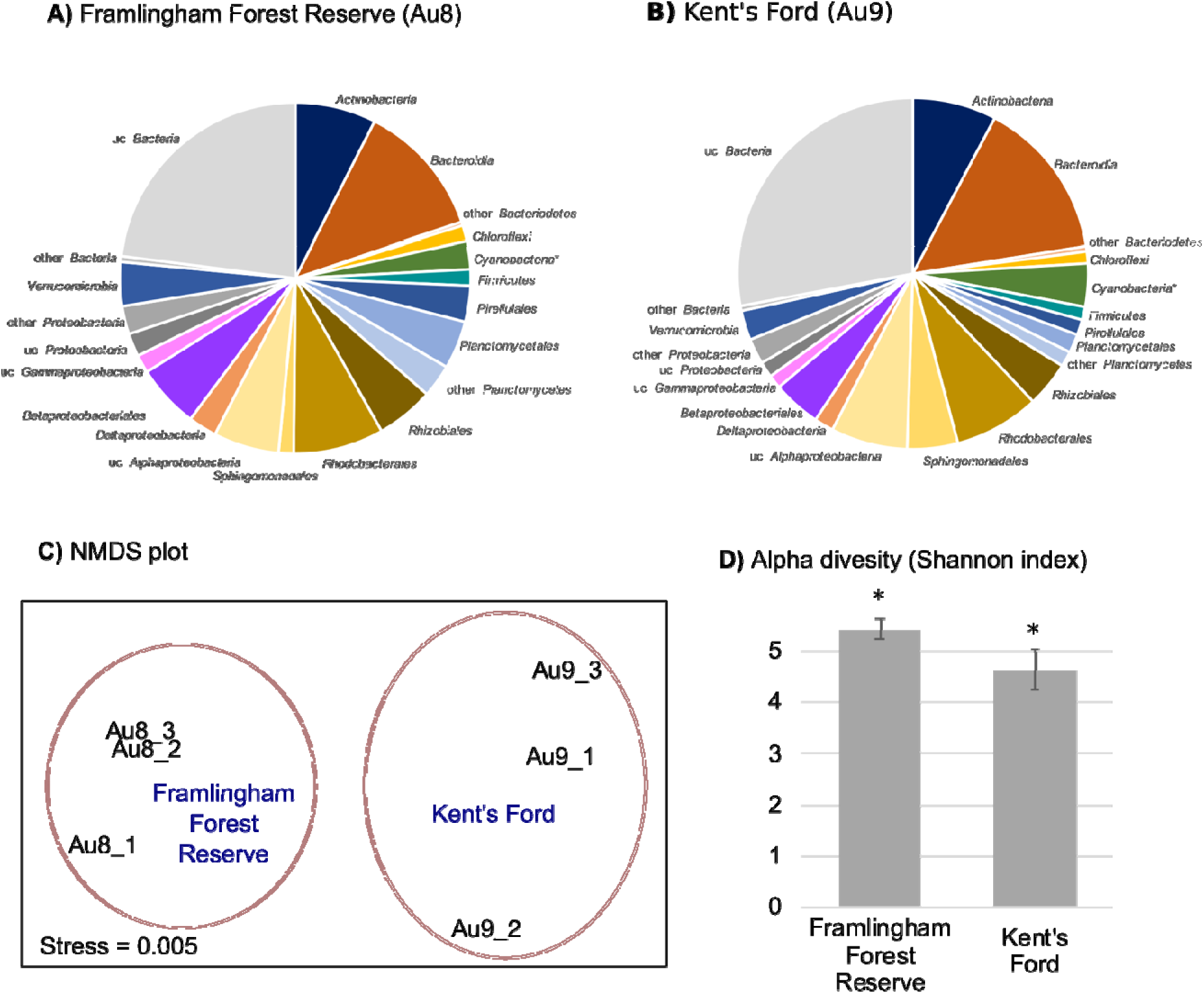
Bacterial community at isolation sites determined by 16S rDNA metabarcoding. Panels A and B show the mean taxonomic distribution of the bacterial communities associated with the triplicate samples at the two sampling sites with sufficient material. Panel C shows the distances between the communities (NMDS plot based on Bray-Curtis dissimilarity matrix); communities at both sites differed significantly (ANOSIM p=0.001). Panel D show the alpha-diversity of bacterial communities in the two sites (mean of three replicates ± SD; * indicates a significant difference, two-sided t-test, p<0.05).

**Figure 5:**
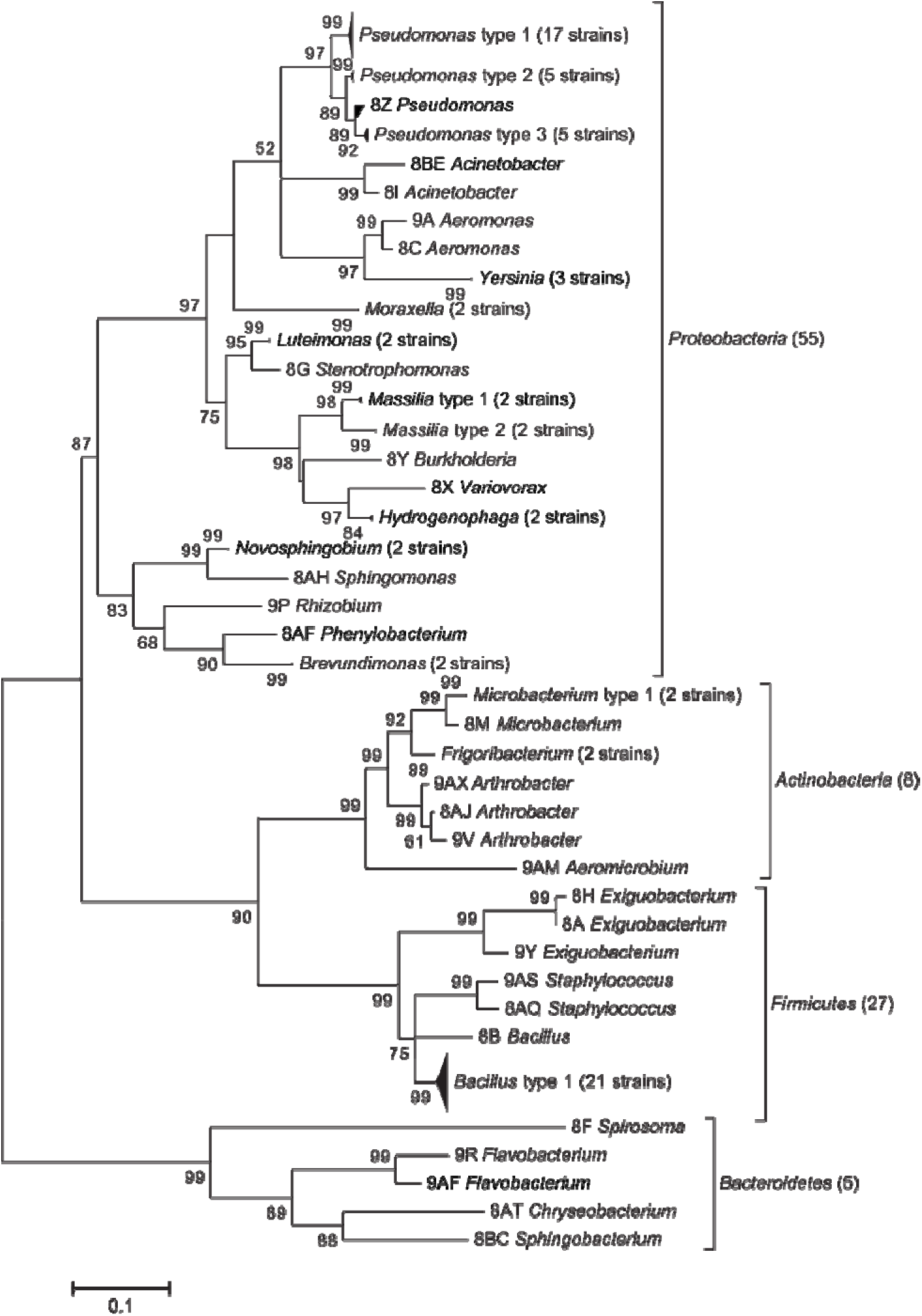
Maximum-likelihood of bacterial isolates obtained from *E. subulatus* in situ. The tree is based on an alignment of 16 rRNA gene sequences of all isolated strain and comprised 598 bases after cleaning. Support values correspond to the percentage of support using 1000 bootstrap replicates. Sequence accession numbers are LR735444-LR735537.

## Discussion

The data presented in this paper provide new information on the distribution of *E. subulatus* in the Hopkins River as well as on its associated microbiota. It confirms that the original finding of *E. subulatus* by West and Kraft was not the result of a transient “contamination”, but that the same population (identical ITS sequence) has likely persisted in the Hopkins River for over 20 years. *E. subulatus* has thus been able to maintain a population despite the water currents. *Ectocarpus* spores and gametes are motile, but swimming speeds reported are in the range of 150-270 μm·s^-1^ (Müller 1978), *i.e.* far less than the speed of the current. This implies that the Hopkins River population either (1) does not rely on gamete releases for reproduction, (2) that its gametes are able to remain close to the substratum as has been suggested for male gametes (Müller 1978) and direct their movement upstream, or (3) that gametes rely on zoochory, as has been proposed in the case of red algae (Žuljević *et al.* 2016). Our findings thus open interesting perspectives for population genetics studies as well as more detailed studies of the reproductive biology of *Ectocarpus* in this area. Furthermore, the fact that no traces of *E. subulatus* were found in nearby rivers or along the coastline suggests that the *E. subulatus* population may be restricted to the Hopkins River, although the range of colonization within the river may have been subject to variation, notably because individuals of *E. subulatus* were no longer found at the original isolation site.

Our sampling campaign also provided novel information on the chemical parameters in the Hopkins River. West and Kraft (1996) reported the salinity at Hopkins River Falls at the time of the isolation of the original strain to be at 1 ppt (approx. 1/34^th^ the salinity of average seawater). This, together with our osmolarity measurements ranging from 74-170 mOsmol, *i.e.* approximately 1/14^th^ to 1/6^th^ of the osmolarity of seawater generally estimated at 1000 mOsmol, leads us to classify the water in the Hopkins River at the sites with *E. subulatus* as low salinity brackish water rather than fresh water. The unusually high osmolarity for a river may be one of the factors enabling *E. subulatus* to be competitive in this environment, a hypothesis which is supported by the fact that no individuals were found in the lower portions of the river with lower osmolarity. High osmolarity in our samples also positively correlated (Pearson correlation p<0.001) with sulfate concentrations between 1 and 5 mM - average sulfate concentrations in fresh water are 0.12 mM (vs. 28 on average in the ocean; Wetzel 2001). Sulfated polysaccharides are typical components of the cell walls of marine plants and algae (Popper *et al.* 2011) and require sulfate for their synthesis, but their importance for *Ectocarpus* remains to be explored. In the same vein the question to what extent the low nitrate concentrations at sites with *E. subulatus* are related to the presence of the algae, either as a cause or as an effect, remains open, although it should be noted that a direct metabolomic comparison of *E. subulatus* and the marine *E. siliculosus* revealed markers for high nitrogen status (total amino acids, ratio of glutamine to glutamate) in *E. subulatus* (Dittami *et al.* 2012). Regardless of the physiological implications of the composition of the Hopkins River water, we argue that it may be more appropriate to refer to the *E. subulatus* strains isolated from the Hopkins River as “fluviatile”, *i.e.* “river” strains rather than freshwater strains, despite their capacity to grow in fresh water in laboratory conditions (Dittami *et al.* 2012).

In addition to these facts about the distribution and environment of *E. subulatus*, the present study provides insights into its associated microbiome – a component likely connected to the capacity of this species to grow in low salinity (Dittami *et al.* 2016). The number of OTUs found on *E. subulatus* in our *in vivo* study was one order of magnitude higher than in a previous study of the laboratory strain after 20 years of cultivation (1312 for six samples from two sites vs 84 OTUs for six samples in two conditions) (Dittami *et al.* 2016). Moreover, a direct taxonomic comparison of these two studies at the genus level revealed only 5 genera (*Acinetobacter, Phycisphaera, Maribacter, Marinoscillum*, and *Gaiella*) that were found in both studies. All of them were rare *i.e.* supported by < 0.01% of reads in our study; Supporting Information File S1). Both studies were based on sequencing runs with similar depth and employed similar analysis pipelines, yet many technical factors could contribute to such differences: the sampling protocol, the primers used, library preparation, sequencing platform and chemistry (Illumina Miseq V2 vs V3) etc.. Nevertheless, the profoundness of the observed differences suggests that either the microbiome of *E. subulatus* in the Hopkins River has evolved and diversified over time or that the cultivation of algae in the laboratory has impacted its microbiome, leading to a reduction of diversity and a change in composition. In a context of the development of new laboratory models for the study of marine holobionts (Dittami *et al.* 2019, preprint), a targeted examination of these potential changes, *e.g.* by following the evolution of alga-associated microbiomes in the field as well as over several cultivation cycles may yield important insights on possible limitations of laboratory model systems. If confirmed, such biases would underline the necessity of devising targeted experiments to test the validity of laboratory findings in the field.

The availability of parallel metabarcoding and untargeted cultivation efforts further allows us to identify under-sampled lineages in cultivation experiments. In this study particularly *Planctomycetes* stand out, as they constituted 8% relative abundance of all algae-associated reads and 176 OTUs but did not have a single associated culture. *Planctomycetes* are notoriously difficult to cultivate, partially due to their long doubling time of up to one month. They require low organic content in media, physical separation from fast-growing competitors *e.g.* via dilution to extinction experiments, and they may benefit from the use of fungicides (Lage and Bondoso 2012). In contrast to the present study, previous barcoding data (Dittami *et al.* 2016) on cultivated *E. subulatus* revealed the presence of very few *Planctomycetes* (0.1% of reads), implying that these culturing techniques would need to be put into place using freshly collected material. In contrast the high abundance of *Firmicutes* in the isolation experiments although they account for only 1% of the reads in the metabarcoding may be due to the fact that these bacteria were particularly amenable to the culture condition.

The present study greatly enhances our knowledge about *E. subulatus* from the Hopkins River, and provides a new set of microbes for coculture experiments. This strengthens the use of *E. subulatus* both as a model for the study of acclimation and adaptation to low salinity and of algal-bacterial interactions.

## Acknowledgments

This work was funded partially by ANR project IDEALG (ANR-10-BTBR-04) “Investissements d’Avenir, Biotechnologies-Bioressources”, the European Union’s Horizon 2020 research and innovation Programme under the Marie Sklodowska-Curie grant agreement number 624575 (ALFF), and an internal call for proposals from the UMR8227 (CNRS, Sorbonne University). We thank Cécile Hervé, Amandine Simeon, and Agnieszka P. Lipinska for helpful discussions; Gwenn Tanguy and Erwan Legeay from the GENOMER platform, Roscoff for support during the library construction and sequencing; and the ABIMS platform for providing the computational facilities for the metabarcoding analyses.

## Author’s contributions

SD, AFP, HK, SE, JW, CB planned the study; TC performed nutrient analyses; AP measured sulfate concentrations; BBD performed metabarcoding experiments; SD performed sampling, culturing, and *in silico* analyses, and wrote the manuscript; All authors corrected the manuscript and approved of the final draft.

## Supporting Information

**Supporting Information File S1– Metabarcoding results.**

